# The cell biology and genome of *Stentor pyriformis*, a giant cell that embeds symbiotic algae in a microtubule meshwork

**DOI:** 10.1101/2024.12.19.629502

**Authors:** Vincent Boudreau, Ashley R. Albright, Therese M. Gerbich, Tanner Fadero, Victoria Yan, Ben T. Larson, Aviva Lucas-DeMott, Jay Yung, Solène L.Y. Moulin, Carlos Patiño Descovich, Mark M Slabodnick, Adrien Burlacot, Jeremy R. Wang, Krishna K Niyogi, Wallace F. Marshall

## Abstract

Endosymbiotic events in which an endosymbiont is retained within a cell that remains capable of phagocytosis, a situation known as mixotrophy, provide potentially important clues about the evolutionary origins of eukaryotes, particularly regarding the relative evolutionary sequence of phagocytosis and endosymbiosis. Mixotrophy in ciliates is commonplace, but has been investigated at a cellular and molecular level almost entirely in one organism, *Paramecium bursaria*. Reliance on just one model system makes it difficult to know which cell biological aspects of this system represent general features of ciliate mixotrophy versus accidental features of the specific organism. Here we describe the cell biology and genome of the giant heterotrichous ciliate *Stentor pyriformis*. We show that this giant unicellular organism contains *Chlorella variabilis* as its endosymbiont, that the *Chlorella* can live freely outside the host, that within the host the *Chlorella* cells are docked near the cell surface, surrounded by microtubule “baskets”, and that the photosynthetic efficiency of the *Chlorella* is reduced inside the *Stentor* cell compared to when it is free-living outside the host, with photon energy instead being shunted to non-photochemical quenching. Compared to the non-mixotrophic *Stentor coeruleus*, *S. pyriformis* has several distinct cellular features that may be related to endosymbiosis: the presence of microtubule baskets, which are absent in *S. coeruleus*; positive rather than negative phototaxis, which is likely an adaptation to allow the photosynthetic symbionts access to sufficient light; and a lack of pigment in the host cell, which may be an adaptation to tolerate high light levels. Compared to *P. bursaria*, *S. pyriformis* has several similar cellular features: in both organisms, the symbiont is a strain of *Chlorella variabilis*; the *Chlorella* endosymbiont retains the ability to live freely when separated from the host; and the algal symbionts contained in perialgal vesicles are docked at the cell surface. One potentially informative difference between *P. bursaria* and *S. pyriformis* is that *S. pyriformis* employs a standard genetic code, similar to other *Stentor* species but different from most other ciliates, including *P. bursaria*, which use a non-standard code in which one or more stop-codons are respecified to encode amino acids. This difference in genetic code could serve as a barrier to impede gene transfer from symbiont to host in other ciliates, but this would not be a factor in *S. pyriformis*. A second cell biological difference is that whereas *P. bursaria* performs phototaxis by a kinetic accumulation mechanism, in which swimming is non-directional but cells slow down in regions of higher light intensity, *S. pyriformis* performs directed swimming towards the direction of high light intensity. However, as in *P. bursaria*, phototaxis in *S. pyriformis* requires the presence of the *Chlorella*, implying a potential flow of information from the symbiont to direct the orientation and swimming of the host cell. We propose that *S. pyriformis* will serve as a useful model system for studying the evolution of mixotrophy and endosymbiosis, with unique advantages in terms of size and regenerative ability as well as distinct cellular and genomic features compared with other mixotrophic ciliate models.

## Introduction

The evolution of the eukaryotic cell is a history of one unicellular organism incorporating parts of another to produce a new organism with improved capabilities. Mitochondria and chloroplasts evolved from free-living organisms that became internalized and converted into organelles dependent on the host cell (Sagan 1967; Lopez-Garcia 2015; Martin 2015; Vosseberg 2024). The first eukaryotic common ancestor (FECA) somehow engulfed another organism without digesting it, after which many more steps of gene transfer to the host cell fixed the internalized cell as an organelle (Maciszewski 2019). These gene transfer processes, with their accompanying changes in protein trafficking and targeting, potentially combined with further uptake events, ultimately gave rise to the last eukaryotic common ancestor (LECA) from which all current eukaryotic cells evolved (Richards 2024). There are various scenarios for how the first eukaryotic cells might have formed. One scenario (known as “phagocytosis first”) suggests that the free-living precursor cells that eventually gave rise to mitochondria were ingested by a cell that already contained the necessary phagocytic machinery (Guy and Ettema 2011; Yutin 2009), but instead of being digested, they were maintained as endosymbionts which then gradually became more dependent on the host cell until they could no longer exist as free-living organisms and thus became fixed as organelles. This scenario has been questioned because having a respiratory organelle would allow the plasma membrane to specialize for phagocytosis (Martin 2017). Two alternative scenarios have been proposed: in one case, one cell type ends up inside another due to some catastrophic event, after which the two cell types need to adapt to the new arrangement. In a second alternative scenario, known as “pre-symbiosis” or “symbiosis first” (Speijer 2020), two cell types develop a symbiotic relationship in which they grow together in close proximity (Dey 2016), and then when one of the cell types evolves the ability to engulf the other, the original symbiotic relationship becomes one of endosymbiosis.

In evaluating these scenarios, it is of great interest to know more about how endosymbiosis can occur, and what cellular or genomic adaptations it may require in the host and symbiont. It is impossible to go back in time to observe what took place during the evolution of FECA and LECA. However, one way to gain insights into potential sequences of events and accompanying mechanisms is to examine extant species in which one cell maintains another cell type within it as an endosymbiont, but where the endosymbiont has retained its ability to live independently of the host and has thus not yet become fixed as an organelle.

Given the importance of determining the relation between phagocytic uptake, endosymbiosis, and eventual organelle fixation, a particularly interesting phenomenon that can still be observed in extant organisms is mixotrophy, in which a phagocytic host cell acquires endosymbionts and maintains them inside itself, while still retaining the ability to capture prey via phagocytosis (Esteban 2010). Such mixotrophic organisms can take up new symbionts by phagocytosis without digesting them, instead maintaining them as endosymbionts. In many cases, the ingested organism is retained fully intact inside perialgal vesicles, while in other cases the plastids are extracted and retained by the mixotrophic host (Johnson 2011). Successful endosymbiosis of intact algal cells inside a phagocytic host requires that the endosymbiont escape digestion and coordinate its division and metabolism with that of the host cell. Meanwhile, the host cell is still capturing prey by phagocytosis, thereby supplementing not only its own nutritional requirements, but also those of the endosymbionts.

The cell biology of mixotrophy raises many interesting questions about how one cell interacts stably with another, for example: what structural adaptations might the host evolve to better maintain the endosymbiont; does the endosymbiont alter its metabolic functions in a way that would benefit the host; and, to what extent does information flow from the symbiont to the host to direct its behavior? Understanding how the host and symbiont cells change their structure and function to work together may provide clues about the structural and regulatory plasticity of cells in general, and may also offer a glimpse into what may have been an important intermediate stage of organelle fixation.

Mixotrophic endosymbiosis with algal cells is seen in many ciliates (Dolan 1992; Esteban 2010), the most classical example being *Paramecium bursaria* (Kodama 2010; He 2019; Jenkins 2024a), which contains algal cells of the species *Chlorella variabilis* as endosymbionts. As with other ciliates that contain algal cells as endosymbionts, the *Chlorella* cells inside *P. bursaria* retain the ability to grow on their own outside the host, and can be re-introduced to host cells from which algae have been removed, the latter also being able to live freely without their symbionts (Kodama 2010). *Chlorella* enter the host cell via phagocytosis, and end up in perialgal vesicles, which are derived from digestive vacuoles that first surround the algae when they are taken up from the environment (Kodama 2022; Fujishima 2022). While inside the host, the cell walls of *Chlorella* are thinner than when the *Chlorella* are free-living (Higuchi 2018), possibly reflecting the activity of glycan-processing enzymes encoded by the *P. bursaria* genome (Jenkins 2024b). The peri-algal vesicles are associated with the cell cortex, showing close association with mitochondria and trichocysts (Kodama 2023).

*Chlorella* cells can divide while inside the *Paramecium* host, and their division is correlated with the division of the host cell (Kadono 2004; Takahashi 2007). The period when the algal cells proliferate matches a period in *Paramecium* division when cytoplasmic streaming pauses. This streaming is microtubule dependent (Nishihara 1999), and inhibition of streaming using microtubule inhibitors is sufficient to trigger *Chlorella* proliferation (Takahashi 2007), pointing to a role for host microtubules in regulating algal replication.

*P. bursaria* is just one of a large number of mixotrophic ciliates. Many retain their endosymbionts in intact form, some retain just the plastids, and in the case of the marine ciliate *Mesodinium*, the retained plastids can undergo division inside the hosts supported by genetic information from nuclei that were ingested and then maintained along with the chloroplasts from the prey (Moeller and Johnson 2023). The ingestion and maintenance of one organism inside another may seem unusual, but in some fresh water aquatic environments, mixotrophic ciliates constitute the majority of ciliate biomass (Finlay 1998; Woelfl 2002), suggesting that mixotrophy has proven to be a highly successful evolutionary strategy.

The fact that many ciliates retain their algal endosymbionts in an intact form, capable of living outside the host, rather than becoming fixed as organelles, suggests that the endosymbionts are not transferring critical genes to the host nucleus. Although ciliates are in some cases capable of horizontal gene transfer (Archibald 2008; Ricard 2006; Shaw 2010; Feng 2020), in species like *P. bursaria* this does not seem to have taken place from the *Chlorella* genome, as judged by the fact that the endosymbionts are still capable of living outside the host and therefore have not lost critical genes (Kodama 2010). Why haven’t these algae become fixed as organelles?

One potential hypothesis is that because most ciliates use a non-standard genetic code in which one or more stop codons specify amino acids (reviewed in Koonin 2017), for example UGA encoding tryptophan, gene transfer from an endosymbiont using a standard code to a host nucleus using an alternative code might be more difficult to achieve in the ciliates compared with other phyla. The recent assembly of the *Paramecium bursaria* genome confirms that it employs a non-standard genetic code and small introns, with increased intron retention of large introns suggesting a bias against correct processing for introns of more standard length (Leonard 2024). On the other hand, there is evidence that *P. bursaria* or its ancestor has undergone horizontal gene transfer from prokaryotes (Jenkins 2024b) so there does not seem to be an insurmountable obstacle to such transfer.

Currently, *P. bursaria* is by far the most extensively studied mixotrophic ciliate at both the cell biological and genomic levels. The fact that most of what we know about cellular and genomic adaptations to mixotrophy in ciliates comes from a single organism makes it hard to know which of the features outlined above may be general features important for mixotrophic interactions with *Chlorella* and which may be mere historical accidents acquired in the lineage of this one organism. By exploring the cell biology of endosymbiosis in other, more distantly related ciliates, in which the acquisition of *Chlorella* endosymbionts is likely to have occurred independently, it would become possible to ask whether any of the specific cell biological features of host-symbiont interaction seen in *P. bursaria* are employed in other independent cases, which would suggest that endosymbiosis may have relied on a common underlying capability shared among ciliates.

Ciliates are classified into two major groups, Postciliodesmatophora and Intramacronucleata (Lynn 2010*). P. bursaria* and indeed most of the mixotrophic ciliate model systems that have been studied, such as *Euplotes*, fall into the Intramacronucleata sub-phylum. Postciliodesmatophora consists of two classes – Karyorelictea and Heterotrichea, the latter being distinguished by ultrastructural features and an unusually large cell size. This group includes the genus *Stentor* (Tartar 1961), famous for their remarkable abilities to heal wounds and regenerate (Marshall 2021), as well as *Spirostomum* (Zhang 2023) and *Blepharisma* (Singh 2023). All of the Intramacronucleata, as well as Blepharisma, use a non-standard genetic code. In contrast, the first *Stentor* species whose genome was determined, *S. coeruleus*, uses a standard genetic code in which all three stop codons are used as stop codons (Slabodnick 2017).

Here, we describe the cellular organization, behavior, and genome of *Stentor pyriformis*, a giant heterotrichous ciliate that, like *P. bursaria*, also hosts *Chlorella variabilis* endosymbionts (Hoshina 2021). There are several reasons for wanting to investigate endosymbiosis in this organism. First, as argued above, understanding the cellular features that enable mixotrophy in a ciliate more distantly related to *P. bursaria* may allow general and essential features to be distinguished from historical accidents. Second, *S. pyriformis* provides an opportunity to examine mixotrophy in ciliates with a standard genetic code. Finally, *S. pyriformis* offers some specific experimental opportunities due to their large size, robust wound-healing, and regeneration capacity, similar to other S*tentor* species, which opens the possibility of microinjection and microsurgical techniques to manipulate the host cell and its symbionts, as well as their tractability for measurement using spectroscopic assays for photosynthetic efficiency and the ease of determining cell orientation during phototaxis.

Our results show that *S. pyriformis* cells construct a system of microtubule structures to hold the *Chlorella* at the cell cortex; that photosynthetic efficiency of the *Chlorella* is reduced when inside the host, possibly playing a role in light protection; that phototaxis in *S. pyriformis* occurs via directional swimming that requires the presence of photosynthetically active endosymbionts, suggesting an exchange of information between the two; and that the macronuclear genome of *S. pyriformis* uses a standard genetic code and has a substantially smaller genome than *Stentor coeruleus*, suggesting it might not have undergone the whole-genome duplications seen in other members of the *Stentor* genus. Taken together, our results show that some aspects of the symbiosis in *S. pyriformis,* such as the type of endosymbiont *Chlorella variabilis*) and docking of endosymbionts near the cell surface, are highly similar to those in *P. bursaria*, while other aspects, such as the standard genetic code of the host and the use of directional phototaxis versus kinetic accumulation, are notably different, confirming that the use of *S. pyriformis* as a model system can lead to new questions about host-symbiont interactions at the cellular level that may be significant for the initial, and still ongoing, evolution of eukaryotic cells.

## Results

### Cell anatomy of Stentor pyriformis

We identified cells of *Stentor pyriformis* in a lake in Falmouth, Massachusetts, based initially on the appearance of the cells as large (300 μm diameter) green trumpet-shaped cells (**Figure 1A; Supplemental Video S1**), sometimes swimming near the surface and sometimes attached to plants in shallow water. Like other *Stentor* species, one end of the cell contains a large membranellar band consisting of thousands of cilia (**Figure 1B**). The cells were covered in longitudinal bundles of microtubules (**Figure 1B**) corresponding to the microtubule bundles known as Km fibers in other *Stentor* species, and contained roughly 5000 green algal cells, based on an automated cell counter measurement of algae isolated from 5 cells. The algal cells have diameters of approximately 3 μm and are mostly located near the cell surface (**Figure 1C**).

**Figure 1.**
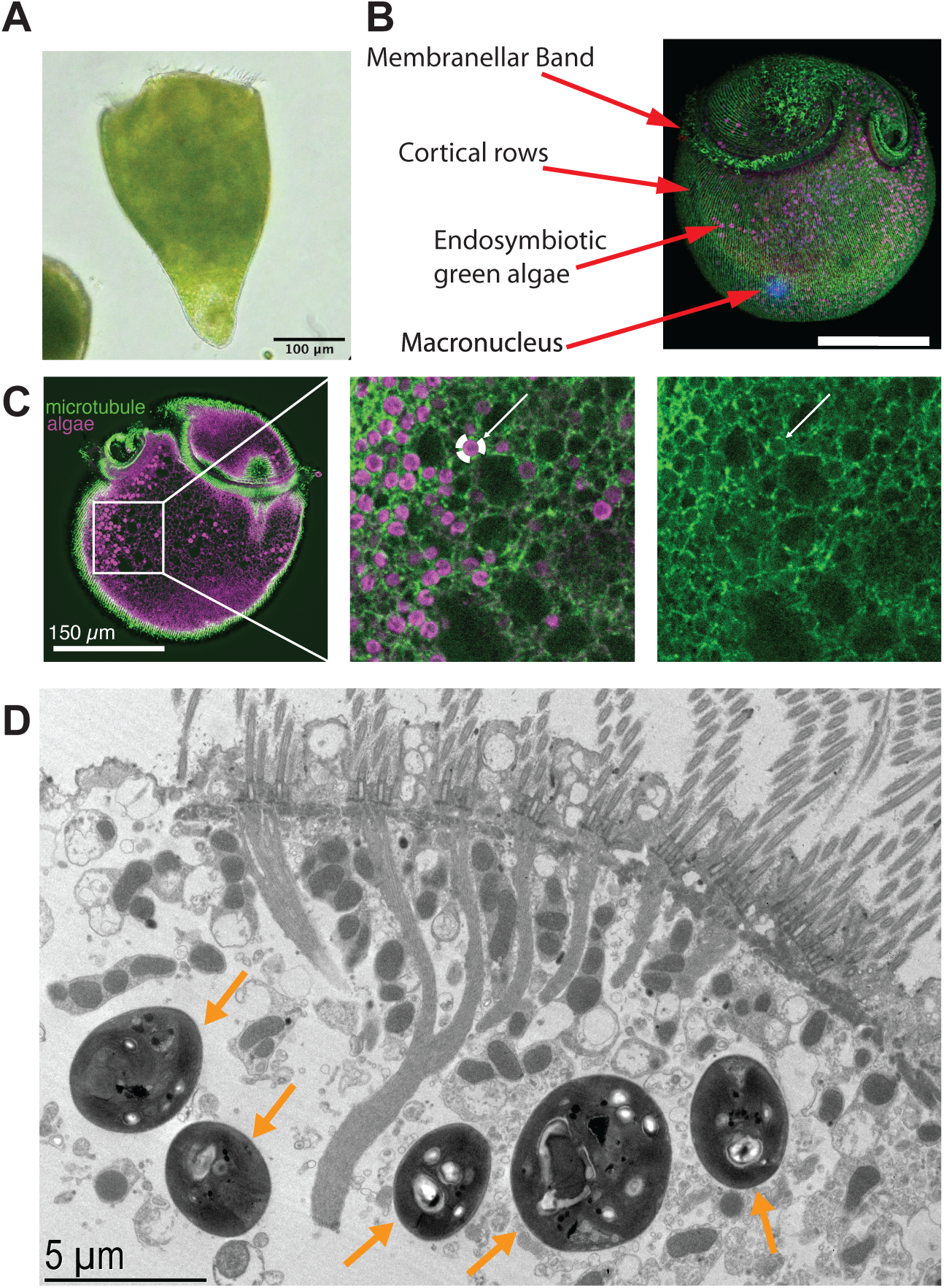
Anatomy of *Stentor pyriformis*. (**A**) Live *Stentor pyriformis* cell imaged by transmitted light microscopy. (**B**) Immunofluorescence image of *Stentor pyriformis*, (green) tubulin immunofluorescence, (pink) algal chlorophyll autofluorescence. Scale bar 250 μm. (**C**) Section through 3D immunofluorescence image showing detail of algal cells docked near cell cortex surrounded by microtubules. (**D**) Thin section electron micrograph of *S. pyriformis* showing algal endosymbionts (orange arrows) in association with cortical structures. Host cell cilia are visible in the upper right corner, and the branching structure is the cortical rootlet system of the membranellar band.

DAPI staining revealed a macronucleus organized as a small number of spherical nodes, often as few as one or two, which is distinct from the moniliform macronucleus seen in *Stentor coeruleus* (Tartar 1961; Mcgillivary 2023) but is in line with the earliest reports of *S. pyriformis* structure which reported a macronucleus containing two large, spatially separated nodes (Johnson 1893) as well as a more recent report on *S. pyriformis* (Hoshina 2021). Tubulin immunofluorescence revealed a dense meshwork of microtubules surrounding the algal endosymbionts at the cell surface, creating an impression of microtubule-based “baskets” surrounding each endosymbiont (**Figure 1C**).

At the ultrastructural level, thin-section electron microscopy of chemically fixed cells showed that the algae inside perialgal vacuoles were interspersed with mitochondria at the cell surface (**Figure 1D**). The endosymbionts contained a prominent pyrenoid with thylakoid membranes penetrating it, as previously reported in *S. pyriformis* isolated from highland bogs in Japan (Hoshina 2021).

### Host and symbiont phylogeny in Stentor pyriformis

For the vast majority of mixotrophic ciliates that contain green algal endosymbionts, the symbiont is a species of *Chlorella*. However, different freshwater ciliates can contain different species of *Chlorella* endosymbionts (Summerer 2008). A study of *S. pyriformis* cells isolated in Japan identified the endosymbiont as *Chlorella variabilis* (Hoshina 2021). We sequenced PCR-amplified 18S rDNA and found that the endosymbiont in our cells isolated in Falmouth, MA are also *Chlorella variabilis* (**Figure 2A**). We confirmed this identification by sequencing a PCR-amplified region of the chloroplast-encoded *rbcL* gene (**Figure 2B**). The *Chlorella* cells can be readily obtained by lysing *S. pyriformis* cells using a glass rod and then spreading the contents of the cell on a MBBM agar plate, after which the *Chlorella* can be grown, independently of the host, on MBBM media (**Figure 2C**).

**Figure 2.**
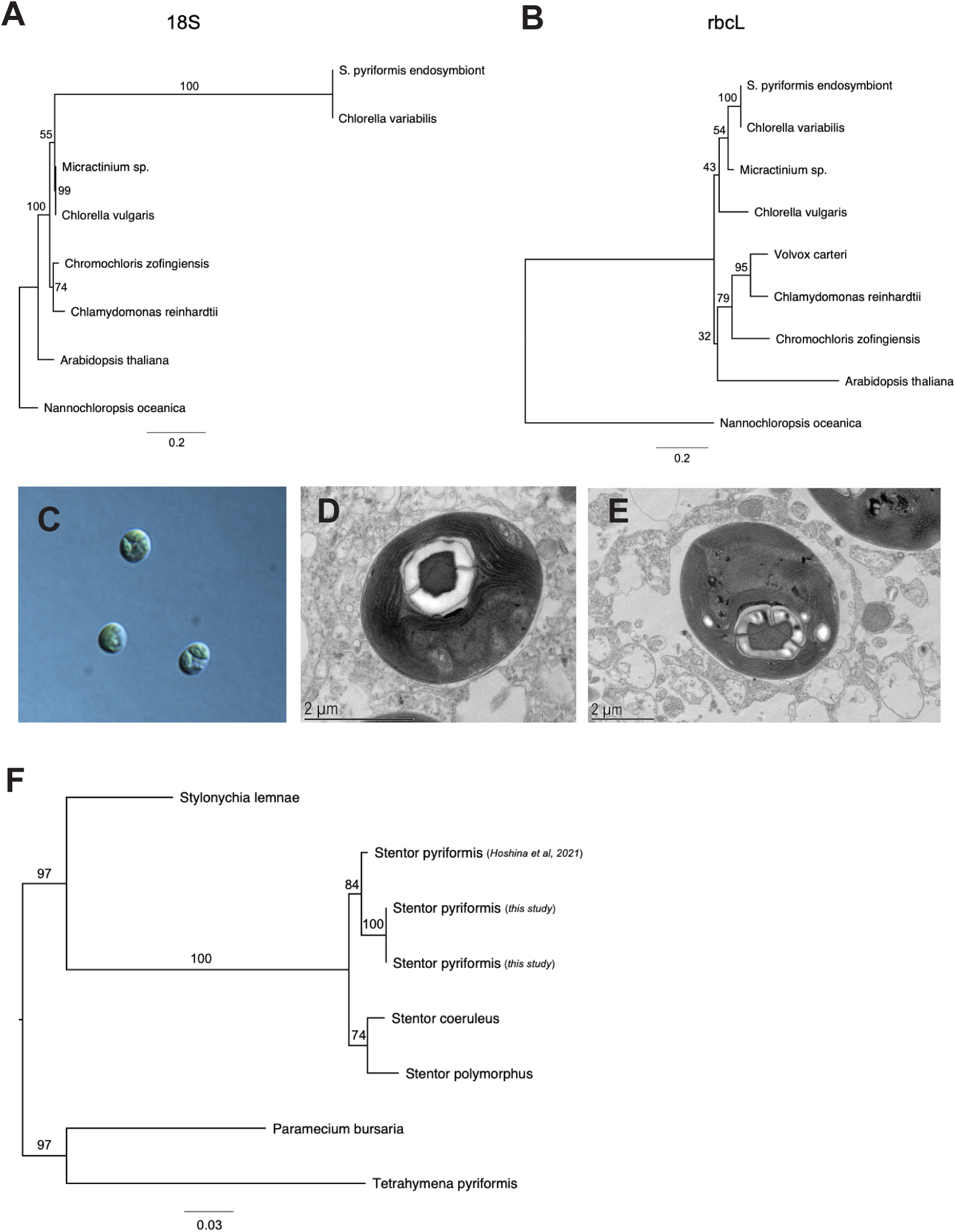
Characterization of the endosymbiont in S. pyriformis. (**A**) Analysis of 18S rDNA sequence indicates that endosymbiont is *Chlorella variabilis*. (**B**) Identity of symbiont confirmed by sequence of the rbcL gene. (**C**) Image of live *Chlorella* cells isolated from *S. pyriformis*. (**D,E**) Thin section electron micrographs showing detail of *Chlorella* ultrastructure. (**F**) Analysis of 18S rDNA sequence confirms that the cells isolated from Falmouth, MA, are indeed *S. pyriformis*.

In *P. bursaria*, which also contains *Chlorella variabilis*, the endosymbionts have cell walls that are approximately half as thick as when they are in the free living state (Higuchi 2018). Our thin-section transmission electron micrographs of chemically fixed *S. pyriformis* cells indicate that the cell wall in the *Chlorella* inside *S. pyriformis* are approximately 30-40 nm thick (**Figure 2D,E**), which is comparable to the thickness of cell walls seen in *Chlorella* inside *P. bursaria* (Kodama 2010).

We initially identified the organism that we collected as *S. pyriformis* based on its size, shape, and presence of green endosymbionts, all of which match historical and modern descriptions (Johnson 1893; Hoshina 2021). In parallel with phylogenetic analysis of the Chlorella endosymbionts, we carried out 18S analysis of the host, which confirmed that we are in fact working with *S. pyriformis* (**Figure 2F**).

### Microtubule baskets surround algal vesicles at the cell cortex in S. pyriformis

Immunofluorescence images show that algal cells at the cell cortex are surrounded by a meshwork of microtubules that extends approximately 3-4 μm into the cell, which is comparable to the diameter of the *Chlorella* cells (**Figure 3A,B).** Counting *Chlorella* cells in sections from 3D images indicates that more than 90% of algal cells are associated with microtubule “baskets”. We obtained higher-resolution images of the microtubule baskets using expansion microscopy (**Figure 3C; Supplemental Video S2**). These images show that the “baskets” do not have any obvious regular structure but appear more like a meshwork of microtubules with gaps where the algae are located.

**Figure 3.**
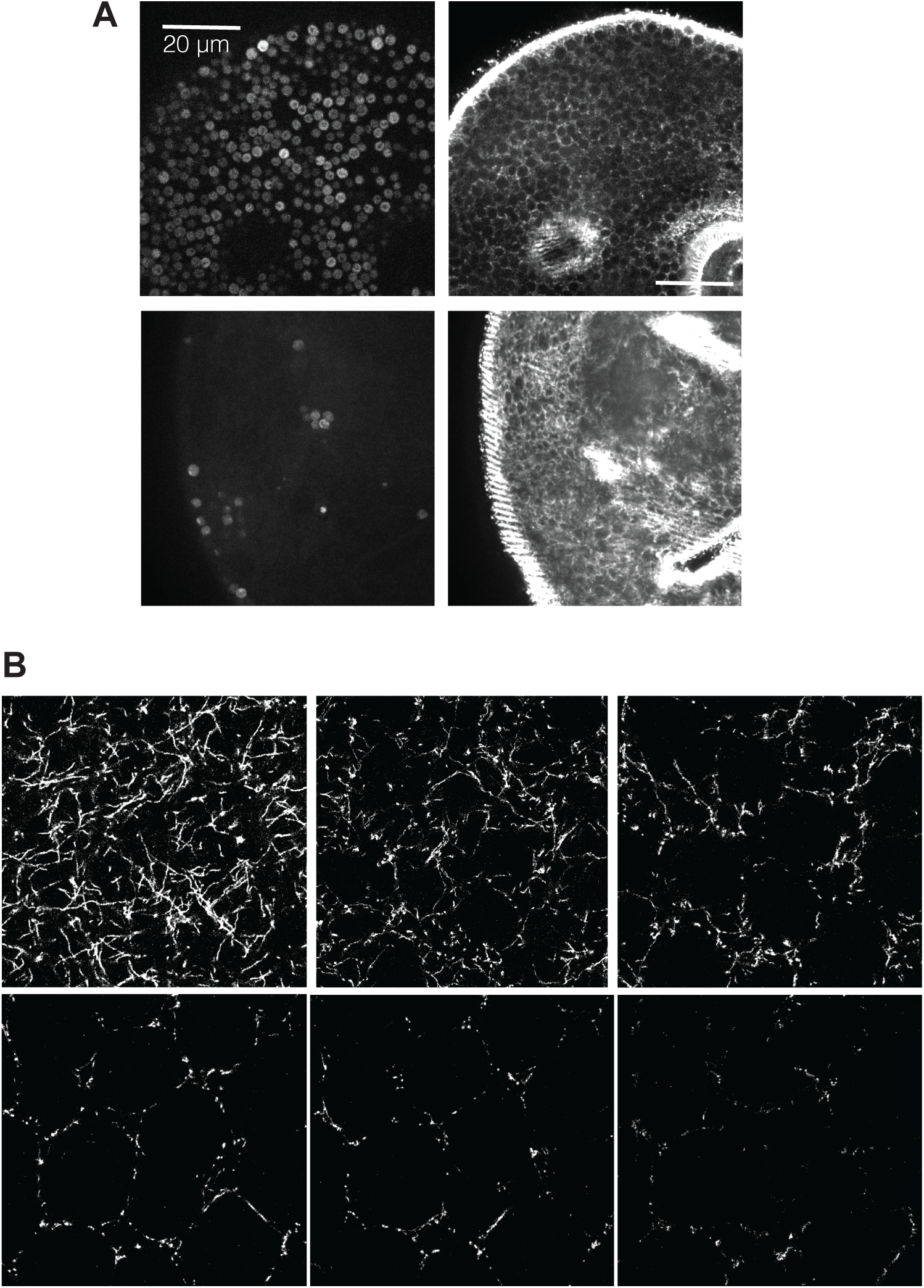
Visualization of microtubule baskets on the host cortex surrounding algal symbionts. (**A**) Chlorophyll autofluorescence of algae (left panels) with Immunofluorescent imaging of beta tubulin (right panels) showing *Chlorella* enriched near the cell cortex in a region containing dense microtubules. (**B**) Expansion microscopy imaging of microtubule baskets. Images show six sequential sections starting from the cell cortex and proceeding inward in 10 micron steps.

The interaction of the endosymbionts in their perialgal vacuoles with the microtubule basket system is highly dynamic. In live images, the symbionts can be seen to move over the surface (**Supplemental Video S3**), and they sometimes disappear from surface patches, suggesting a continuous remodeling of the microtubule network. When cells are centrifuged at 4000 g, the algal cells detach from the cortex and pile up on one side of the cell. After such a perturbation, the algae return to the cortex on the time scale of tens of minutes, again consistent with a dynamic interaction with the cortical microtubule network. A similar ability of *Chlorella* endosymbionts to be displaced by centrifugation and then re-attach to the cortex has been shown in *P. bursaria* (Kodama and Fujishima 2013; 2023).

### Clearing of Chlorella endosymbionts from the Stentor host

In order to obtain *Stentor* host cells that were free of endosymbionts, we developed a protocol in which cells are centrifuged at 10000 g, which causes the alga to accumulate on one side of the cell (**Figure 4A**). Upon centrifugation for 1 minute, cells burst and release their algal contents. Due to the well-known wound-healing ability of *Stentor* cells (Marshall 2021), the host cells are not killed by this procedure, which results in actively swimming alga-free host cells. As an alternative approach, it is also possible to bisect cells with a glass needle after the algae have been collected on one side by centrifugation (**Figure 4B**). The resulting alga-free cells are white in color, thus apparently lacking the blue or purple pigments found in pigmented *Stentor* species like *S. coeruleus* or *S. amethystinus*. In other *Stentor* species (Yang 1986) as well as other pigmented ciliates (Finlay and Fenchel 1986), the pigments are thought to produce reactive oxygen in the presence of light. The absence of pigment in *S. pyriformis* host cells may thus be related to their need to expose themselves to sunlight in order to allow effective photosynthesis by the endosymbionts.

**Figure 4.**
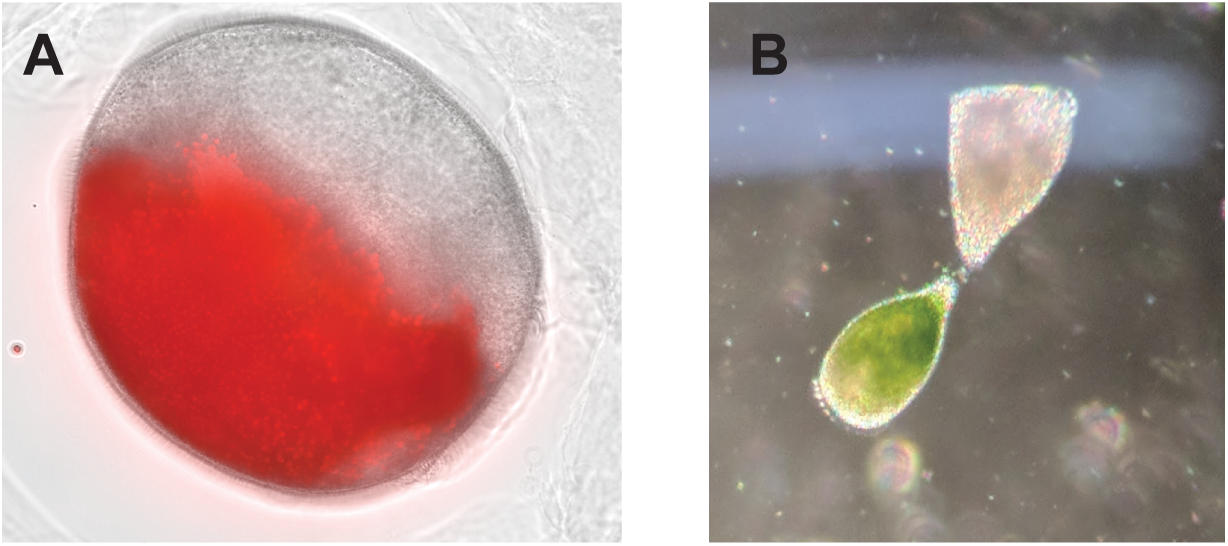
Separating host from symbiont. (**A**) Accumulation of algal endosymbionts on one side of a centrifuged cell as visualized by chlorophyll autofluorescence (red). (**B**) Example of a cell cleared of algae with a normal cell for comparison.

Cells from which the algae have been removed continue to undergo swimming motion for at least a few hours, but they do not undergo division and eventually die. Our experiments do not rule out the possibility that symbiont-free host cells could be kept alive with appropriate nutritional support.

### Reduced photosynthetic efficiency in Chlorella endosymbiont when inside the Stentor host

One of the key questions with respect to endosymbiosis is how, or whether, both the host and the symbiont benefit from their mutual interaction. With this idea in mind, we asked whether the *Chlorella* inside of *S. pyriformis* might show altered photosynthetic efficiency. When a photon is absorbed by chlorophyll, the energy can be used for photochemistry, in which the energy from the photon is (1) used to transfer an electron from the p680 reaction center chlorophyll of photosystem II (PSII) to the primary quinone Q_A_, (2) released as heat (non-photochemical quenching, NPQ), or (3) emitted via fluorescence (Müller 2001). Because fluorescence competes with photosynthesis, it can be used as a measure of photosynthetic efficiency (Baker 2008).

Pulse-Amplitude Modulated (PAM) chlorophyll fluorometry is a common and non-invasive method for determining photosynthetic parameters based on chlorophyll fluorescence (Brooks and Niyogi, 2011). By using saturating and photosynthesis-activating actinic light pulses, chlorophyll fluorescence can be induced and quenched. Of critical importance, photosynthetic efficiency can be determined by measuring photosystem II quantum efficiency, while non-photochemical quenching (NPQ) can be calculated simultaneously using PAM fluorometry.

The results of our analysis show that the photosystem II efficiency (<λPSII) of the *Chlorella* cells inside the host cell is ∼30% lower than that of algal cells cultured outside the host (**Figure 5A**). In contrast, our PAM measurements showed a dramatic increase in their capacity for NPQ when the *Chlorella* are inside the host compared to when they are free-living (**Figure 5B**). One key form of NPQ is energy-dependent quenching, denoted qE, which is a mechanism triggered by high proton concentrations in the thylakoid lumen of the chloroplast. The qE dissipates absorbed light energy from the PSII light-harvesting antenna in the form of heat (Müller 2001). By collapsing the proton gradient across the thylakoid membrane, the proton and potassium ionophore nigericin blocks qE (Brooks 2013). PAM measurements in *S. pyriformis* show that NPQ was blocked by the addition of 15 μM nigericin (**Figure 5B**), suggesting that the NPQ largely consists of qE.

**Figure 5.**
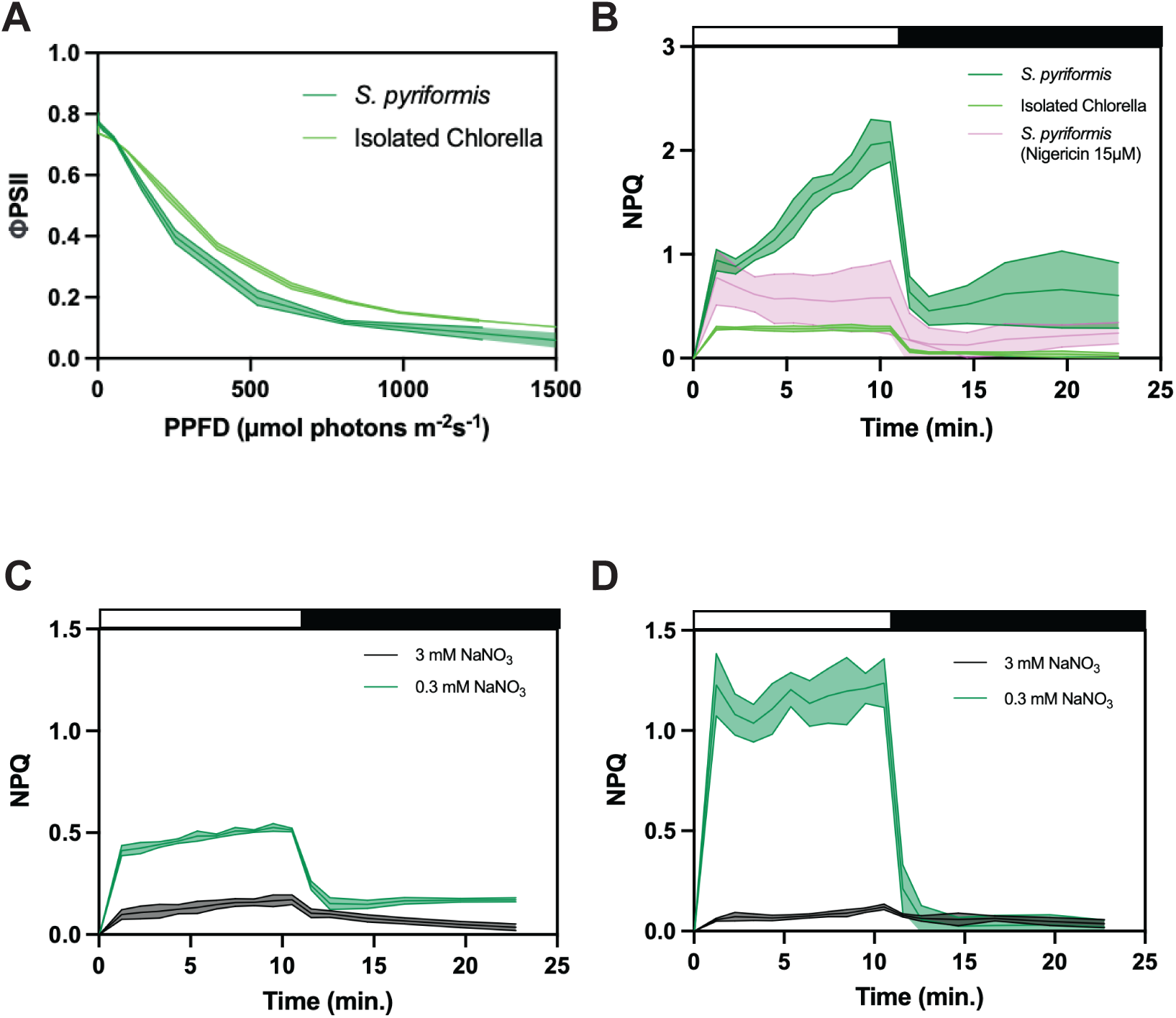
Measurement of photosynthetic efficiency of *Chlorella* cells within *Stentor pyriformis* using PAM fluorometry. (**A**) Photosystem II operating efficiency (<λPSII) as a function of photosynthetic photon flux density (PPFD). (**B**) NPQ as a function of time during and after exposure to actinic light. (**C**) Increased NPQ in free-living *Chlorella* grown with low nitrogen (0.3 vs. 3 mM NaNO_3_) under ambient CO_2_. (**D**) Further increase in NPQ in free-living *Chlorella* grown with low nitrogen under high CO_2_ (3%) conditions. In panels B-D, the white bar above the graph represents exposure to actinic light, and the black bar represents recovery in the dark.

Symbioses involving *Chlorella* are typically nitrogen-limited, which is thought to control the proliferation of the symbiont (Reisser 1976; McAuley 1996; Esteban 2010). Consistent with such a situation in *S. pyriformis*, the high level of NPQ activity that we have detected in *Chlorella* inside *S. pyriformis* only occurs in free-living *Chlorella* when they are nitrogen limited (**Figure 5C**). The effect is even stronger when low nitrogen is combined with high CO_2_ (**Figure 5D**). Similarly to *S. pyriformis* NPQ, low-nitrogen-stimulated NPQ in free-living *Chlorella* is sensitive to nigericin. Taken together, these experiments suggest that *S. pyriformis* is able to maintain a level of photosynthetic efficiency in its endosymbionts, while activating NPQ mechanisms for photoprotection through nitrogen limitation in its cytoplasm.

### Phototaxis in Stentor pyriformis uses directional swimming and requires endosymbionts

We observed that *S. pyriformis* cells tended to accumulate in containers on the side facing the nearest light source, suggesting phototactic behavior. *Stentor coeruleus*, a *Stentor* species without symbiotic algae, has been shown to exhibit photophobic swimming behavior (negative phototaxis) when exposed to light (Kim et al., 1984). However, when we placed *S. pyriformis* cells into an experimental phototaxis assay (**Figure 6A**), we found that, similar to other ciliates containing algal endosymbionts such as *P. bursaria* (Matsuoka and Nakaoka, 1988), *S. pyriformis* accumulates near a light source. However, unlike *P. bursaria,* which shows a photo-accumulation behavior in which cells move more slowly in brighter regions so that they gradually accumulate there, *S. pyriformis* swims directly towards light (**Supplemental Video S4**). During this phototactic motion, we see that the cells clearly orient with their anterior ends towards the light source, and they then swim persistently in that direction.

**Figure 6.**
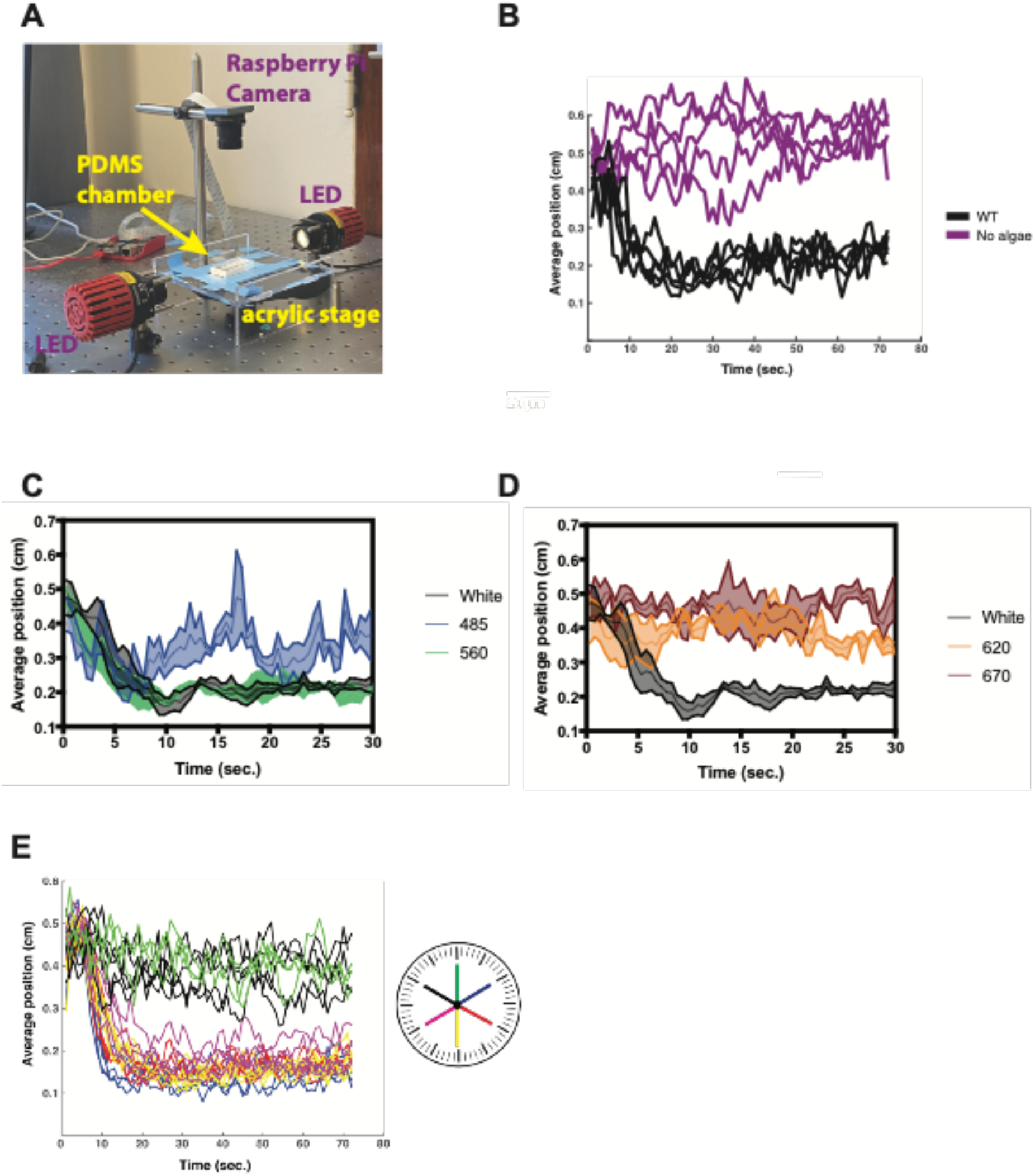
Phototaxis in *Stentor pyriformis*. (**A**) Apparatus for measuring phototaxis. (**B**) Phototaxis measured by increased accumulation of cells at one end of the chamber. (black) untreated cells showing phototaxis. (purple) cells from which algae were removed, showing loss of phototaxis. (**C,D**) Phototaxis measured using light of different wavelengths, showing maximal phototaxis for light at 620 nm. (**E**) Circadian variation in phototaxis. Color code indicates time of day. Cells in these experiments were grown in constant light for a minimum of two days prior to the measurement.

The motile force for phototaxis is provided by the cilia of the *Stentor* host cells, but which organism, the host or the symbiont, is responsible for sensing the light? Given that non-symbiotic *Stentor* species can also detect light (Kim 1984), one possibility is that *S. pyriformis* uses a similar phototaxis system, but with different regulatory connections that drive positive rather than negative phototaxis such as is seen in non-photosynthetic *Stentor* species. Alternatively, it is possible that the algal endosymbionts, which can also detect light, might be serving as the primary photosensors and somehow directing the host cell to swim towards increasing light intensity. To test this idea, *S. pyriformis* host cells were cleared of algae as described in **Figure 4**, and then tested for phototaxis ability. We found that such alga-free cells no longer undergo phototaxis (**Figure 6B**), indicating that the presence of algal endosymbionts is required for phototactic motility of the host cell.

To further characterize phototaxis in *S. pyriformis,* we examined its swimming behavior to different wavelengths using illumination filters (**Figure 6C,D**). All land plants and green algae (Chloroplastida) use chlorophylls a and b as light-absorbing molecules in photosynthesis (reviewed in Bhattacharya et al., 1998). Both chlorophylls a and b absorb blue (∼425-475 nm) and red (∼625-675 nm) light. Interestingly, *S. pyriformis* does not swim at all towards red or orange light (670 or 620 nm) and shows reduced phototaxis towards blue light (485 nm).

Instead, *S. pyriformis* prefers swimming towards green light (560 nm). Although a *Stentor*-based green light photoreceptor cannot be ruled out, several green light photoreceptors have been identified in green algae (Nagel et al., 2002).

Finally, we note that phototaxis in *S. pyriformis* obeys a circadian rhythm (**Figure 6E**) similar to the unicellular alga *Chlamydomonas reinhardtii* (Johnson et al., 1991).

### The draft genome of Stentor pyriformis

We assembled the macronuclear genome of *S. pyriformis* using total genomic DNA as described in Materials and Methods. The current assembly is 29.2 Mb of sequence distributed over 203 contigs with 15,631 predicted genes. We identified telomere sequences using Tandem Repeats Finder (Benson 1999) and filtered repeats for all possible permutations of the 8-mer ‘CCCTAACA.’

We found that 89 contigs have 2 telomeres, 60 have 1 telomere, and 24 have 0 telomeres. In addition, 25 contigs have 3 telomeres, 3 contigs have 4 telomeres, and 2 contigs have 5 telomeres. Additional analyses and curation of the genome are necessary to determine if these contigs should be split at internal telomere repeats or left intact. In particular, we cannot currently determine whether these represent alternative telomere addition sites (ATAS) that may occur in ciliates (Singh 2023). Excluding contigs shorter than 30 kb, the mean contig length is 185 kb and standard deviation is 51 kb. Including shorter contigs that may be removed or combined after further manual curation otherwise artificially lowers the mean and raises the standard deviation of contig length. As shown in **Figure 7A**, this length distribution is substantially narrower than that of a draft long-read assembly of *S. coeruleus* (Albright (Forthcoming 2024). PacBio whole-genome sequencing and draft assembly of Stentor coeruleus [Dataset]. Dryad. https://doi.org/10.5061/dryad.bzkh189kk). We note here that genome comparisons between *S. pyriformis* and *S. coeruleus* are done with the more contiguous draft genome than our published reference genome (Slabodnick 2014). The GC content is comparable to *S. coeruleus* although shifted slightly higher (**Figure 7B**). *Stentor coeruleus* was notable for its use of extremely small introns, almost all of which were 15 nt in length (Slabodnick 2017). To account for short introns in gene prediction, we used Intronarrator (https://github.com/Swart-lab/Intronarrator, Singh 2023), which is a wrapper that predicts and removes short introns, runs Augustus with an intronless model, and adds the introns back in the end (see Methods). We found that while many introns are also short in *S. pyriformis*, there is also a substantial tail of longer introns that was not observed in *S. coeruleus* (**Figure 7C**).

**Figure 7.**
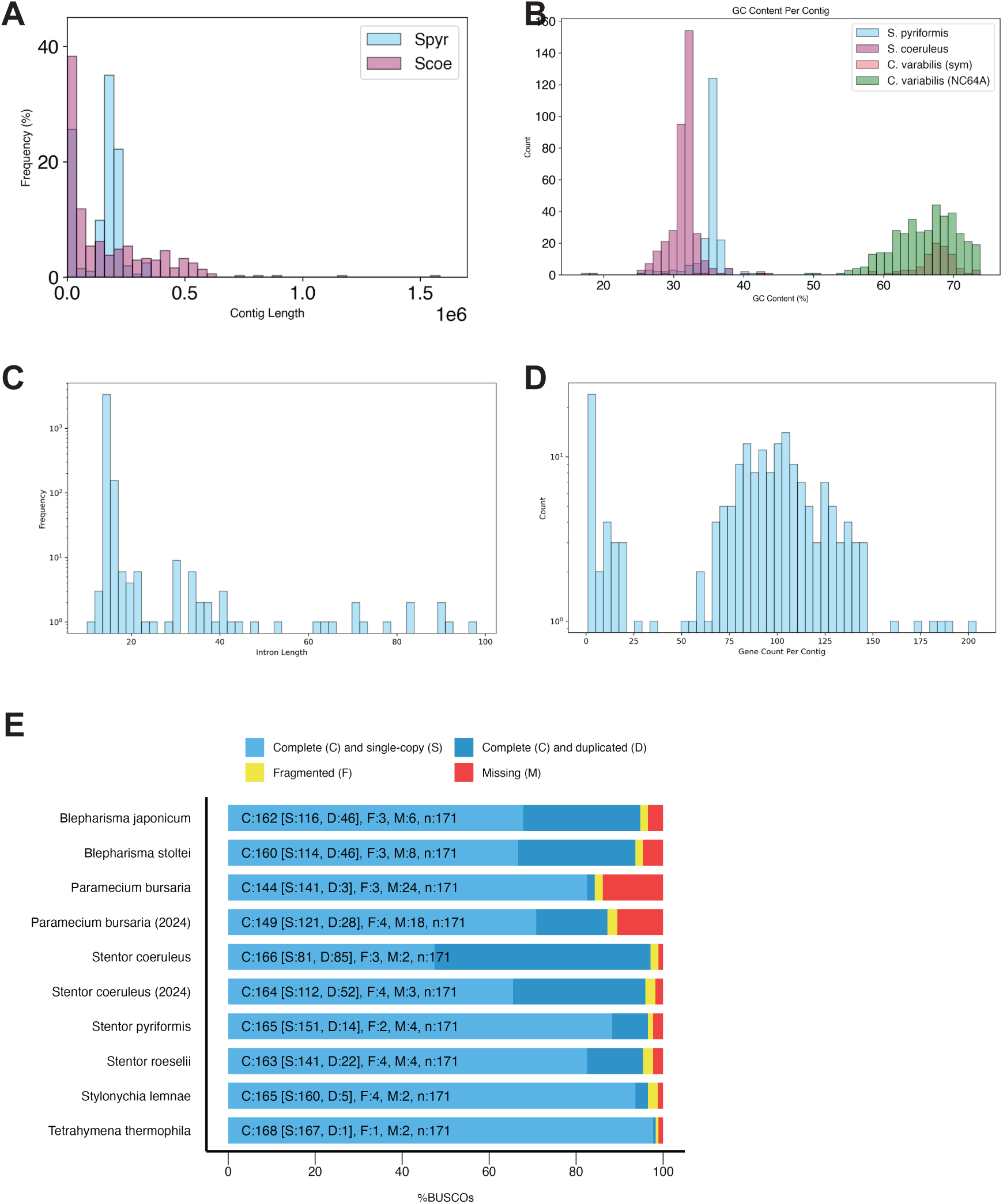
Genome of *Stentor pyriformis*. (**A**) distribution of chromosome (contig) sizes in assembly of *S. pyriformis* versus *S. coeruleus* genomes. (**B**) GC content for the genomes of S. pyriformis (host and symbiont) compared to S. coeruleus and C. variabilis NC64A. (**C**) Distribution of intron lengths. (**D**) Distribution of predicted gene numbers per contig. (**E**) BUSCO analysis of *S. pyriformis*.

We note that the contigs we have identified are assumed to represent only the macronuclear genome, given the much higher ploidy compared with the diploid micronucleus. At this point we cannot strictly rule out the possibility that the assembly might include some micronuclear sequence.

### Stentor pyriformis macronuclear genes use a standard genetic code

One unusual feature of most ciliate genomes is their use of non-standard genetic codes, in which some codons specify different amino acids, or mostly notably, where one or more stop codons no longer serve as stop codons but instead are used to encode amino acids, such as UAA or UAG being translated as glutamine or UGA being translated as cysteine (Tourancheau 1995; Lozupone 2001; Swart 2016; Chen 2023). We previously found that *S. coeruleus* uses a standard genetic code in which all three stop codons are used as stop codons (Slabodnick 2017). To determine if this is also true for *S. pyriformis*, we used Codetta to predict the genetic code (Shulgina 2023). Codetta predictions indicated the use of the standard genetic code; however, Codetta only predicts the genetic code used in coding sequences, and cannot predict stop codon reassignments such as those that are prevalent in ciliates. Thus, we also examined tRNAs predicted by tRNAscan-SE_2.0 (Chan 2021) which revealed tRNAs for all of the standard amino acids as well as selenocysteine. One tRNA was annotated as a suppressor tRNA, with an anticodon complementary to UGA, but the gene encoding this tRNA mapped to the mitochondrial contig. UGA-targeting tRNA is a shared feature of ciliate mitochondrial genomes (Inagaki 1998) and does not affect the stop codon usage of macronuclear genes. Counts of the final three nucleotides for every predicted coding sequence also showed that all three stop codons were used with different frequencies (UAA 62.0%, UAG 30.5%, UGA 7.5%). Therefore all data indicate that *S. pyriformis*, like *S. coeruleus* but unlike *P. bursaria*, uses a standard genetic code.

### Reduced gene complement compared to Stentor coeruleus

The evolution of ciliates has involved a number of whole-genome duplications that help explain the large number of genes found in many ciliates, but it has been noted that in other ciliates with endosymbionts, there are far fewer genes, suggesting less extensive duplication. To ask whether this would also be true in *S. pyriformis*, we predicted gene models from our *S. pyriformis* macronuclear genome assembly as described in Methods. This resulted in approximately 15,000 predicted genes.

Ciliates vary greatly in the organization of macronuclear contigs, with some species having as few as one gene on each contig. Single-gene contigs could potentially have a narrower length distribution than contigs with larger gene numbers. With the predicted gene models in hand, we found that the average predicted number of genes per contig is approximately 100 (**Figure 7D**).

One feature of the *P. bursaria* genome is that specific gene families, notably involved in oxygen binding, are depleted relative to other ciliates (He 2019). One explanation for missing genes in a mixotrophic ciliate could be that the presence of the symbiont relieves the host from requiring genes encoding metabolic activities that the symbiont normally provides. In order to ask whether this might be true in *S. pyriformis*, we carried out a BUSCO (Benchmarking Universal Single-Copy Orthologs) analysis, which is based on a set of highly conserved genes among a wide swath of biodiversity (Manni 2021). BUSCO is primarily intended as a benchmark for genome completeness, but it can also give an indication of genomes that, for reasons such as symbiosis or a parasitic lifestyle, have undergone extensive gene loss. BUSCO analysis of *S. pyriformis* using the alveolata_odb10 database (**Figure 7E**), showed a similar number of complete versus missing genes compared to other *Stentor* species that do not contain *Chlorella* endosymbionts, in contrast to *P. bursaria* which clearly has a substantially larger number of missing genes. Given the relatively high quality of the *P. bursaria* genome, this difference may reflect actual gene loss in *P. bursaria* compared to *S. pyriformis*.

The number of predicted *S. pyriformis* genes, approximately 15,000, is substantially smaller than the 34,506 genes annotated in our non-symbiotic *Stentor* species *S. coeruleus* reference assembly (Slabodnick 2017). While our annotations of the long-read draft *S. coeruleus* assembly are incomplete, we observe a slight reduction in the percentage of duplicated BUSCOs compared to the assembly (**Figure 7E**) and believe the number of *S. pyriformis* genes will remain markedly smaller than *S. coeruleus*.

### The genome of Chlorella variabilis from Stentor pyriformi***s***

We were able to release the *Chlorella* cells into MBBM media (**Figure 2C**) and culture them on MBBM media agar plates (Methods). We sequenced and assembled the genome of the *Chlorella* symbiont (see Methods) using total DNA isolated from cultured *Chlorella* cells derived from *S. pyriformis* cells. The assembled *Chlorella* genome was 49.6 Mb in size, distributed across 93 contigs and scaffolds with 11,914 predicted genes. For comparison, the best characterized *C. variabilis* species (NC64A) isolated from *P. bursaria* had an assembled genome of 46 Mb, with 12 chromosomes and 9,791 predicted genes (Blanc 2010). The GC content of the *Chlorella* from our cells has the same distribution as reported for C. variabilis NC64A (**Figure 7B**). The contig size in our *Chlorella* assembly mostly agrees with that reported for NC64A but has a few larger contigs (**Figure 8A**). BUSCO analysis indicates a similar completeness of our *Chlorella* genome as that of NC64A (**Figure 8B**). We note that the *Chlorella* endosymbiont isolated from *S. pyriformis* in Japan (Hoshina 2021) was also closely related to NC64A. One tends to think of the host as being more complex than its endosymbionts, and in this light it is interesting to note that the genome of the *Chlorella* endosymbiont in *S. pyriformis* is substantially larger than the genome of the host itself.

**Figure 8.**
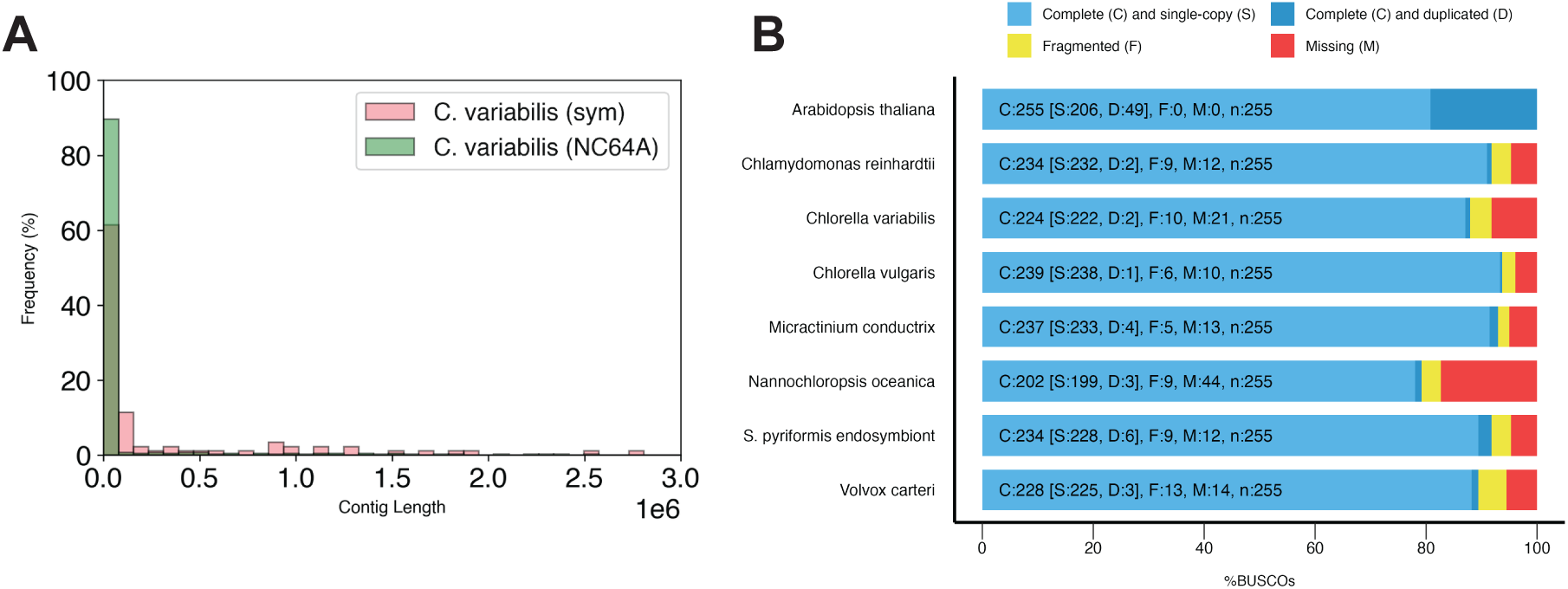
Genome of *Chlorella variabilis* isolated from *Stentor pyriformis*. (**A**) Distribution of chromosome (contig) sizes in assembly of the *Chlorella* endosymbiont isolated from *S. pyriformis* versus *C. variabilis* NC64A. (**B**) BUSCO analysis of *C. variabilis* genes and predicted proteins from our isolate of *S. pyriformis*.

In light of the high NPQ capacity of the *Chlorella* endosymbionts in *S. pyriformis* (**Figure 5B**), we confirmed that the *Chlorella* genome in our isolates encodes orthologs of chlorophycean violaxanthin de-epoxidase (CVDE) (Li et al., 2016 Nature Plants), zeaxanthin epoxidase (ZEP) (Baroli et al., 2003 Plant Cell), and stress-related light-harvesting complex (LHCSR) (Peers et al., 2009 Nature), all of which are involved in NPQ and photoprotection. LHCSR is required for the rapid induction and relaxation of the qE component of NPQ in the light-harvesting antenna. CVDE, present in the green algae on the stromal side of the thylakoid membrane, is involved in synthesis of zeaxanthin in high light for NPQ and reactive oxygen scavenging. ZEP is involved in the removal of zeaxanthin in limiting light. All three proteins show high identity with orthologs in related algae (**Supplemental Figure S1**).

## Discussion

### Implications of the standard genetic code in S. pyriformis for the question of organelle fixation

A notable feature of endosymbiosis in ciliates is the rarity with which endosymbionts become fixed as permanent organelles, with far more cases existing in which the algae retain the ability to live free of the host (Esteban 2010; Johnson 2011). Such fixation is thought to generally require extensive gene transfer from the endosymbiont to the host, such that the symbiont becomes dependent on the host for the expression of one or more essential genes, and that this dependency eventually results in a symbiont that cannot live apart from the host and is thus considered an organelle. One hypothesis for why such fixation is so rare in ciliates is that most ciliates use a non-standard genetic code, in which some of the standard stop codons encode amino acids.. Our analysis of the *S. coeruleus* genome provided extensive demonstration that *S. coeruleus* uses a standard genetic code, with all three stop codons used as stop codons and no tRNA genes with anticodons that could recognize stop codons (Slabodnick 2017). Here, we show that *S. pyriformis* also uses a standard genetic code. Thus, in this particular case, the explanation for why the symbiont has failed to become fixed as an organelle is unlikely to involve the use of a non-standard genetic code by the host.

### Implications of reduced photosynthesis inside the host

The reduced photosynthetic efficiency of *Chlorella* inside the host (**Figure 5**) may suggest that the symbiont is gaining some benefit from being inside *Stentor* that outweighs the reduced ability to photosynthesize, or more simply that its photosynthetic efficiency is limited by available nitrogen. We speculate that possible benefits might include protection from predation due to the huge size of a *Stentor* cell, which makes it almost impossible to be ingested by some filter feeders; nutritional supplementation for the alga from the host which, as a filter feeder itself, may have access to food organisms that provide otherwise scarce nutrients that the *Chlorella* can use for growth; and the swimming ability of the *Stentor* host which allows the otherwise immotile *Chlorella* to move from place to place allowing continued access to light. These hypotheses will require direct testing in future experiments. If we grant that the alga is trading reduced photosynthesis for some benefit obtained from the host, could there be any benefit for the host from reduced photosynthesis? Photosynthetic organisms contain mechanisms to protect themselves and their genomes from photo-oxidative damage due to the sometimes intense sunlight to which they are exposed. Compared to non-symbiotic *Stentor* species, which are best collected in shaded regions of ponds, we routinely find that *S. pyriformis* is abundant near the surface in open regions of water exposed to direct sunlight, either floating near the meniscus or attached to plants near the surface. This spatial distribution is consistent with our observation of positive phototaxis in *S. pyriformis* (**Figure 6**). However, it poses a potential hazard for the host, whose own genome may become exposed to enough light to produce extensive DNA damage. Our PAM measurements of *Chlorella* photosynthetic activity in *S. pyriformis* indicate that the *Chlorella* may be collecting and then shunting energy from absorbed light in such a way as to provide photoprotection. Given the spatial location of endosymbionts near the *Stentor* plasma membrane, we speculate that this might serve to help protect the host genome, in addition to protecting the endosymbiont from excess light. Future work could test this idea by removing the algae, exposing the cells to excess light, and measuring the degree of DNA damage. Such an effect has been seen in *P. bursaria*, in which cells cleared of endosymbionts are substantially more sensitive to UV light than when the symbionts are present (Summerer 2009).

*P. bursaria* has been shown to redistribute its algal endosymbionts when exposed to high light (Summerer et al., 2009). In this case, the algae which are normally located near the cell surface move to a cluster in the posterior region of the cell. Whether this is to provide a denser screen for the benefit of the algae themselves or for host structures like the macronucleus is still uncertain. We have not observed such a rearrangement of endosymbiont position in *S. pyriformis*.

### A model for phototaxis in *S. pyriformis* based on oxygen generation by the endosymbionts

A key question concerning phototaxis of *S. pyriformis* is which organism, the host or endosymbiont, is detecting the light. We have found that phototaxis ceases in cells from which the algae have been surgically removed (**Figure 6B).**

The dependence of phototaxis on the presence of algae is consistent with what has previously been reported in *P. bursaria* and other *Chlorella*-bearing ciliates (Niess 1981; Cronkite 1981; Iwatsuki and Naitoh 1988; Finlay 1987). One difference with *S. pyriformis* is that photo-accumulation of *P. bursaria* occurs equally well in red and in blue-green light (Niess 1981), possibly suggesting a difference in the light-sensing mechanism between the two species. Classical algal phototaxis, such as in *Chlamydomonas*, relies on an eyespot with a fixed directional orientation relative to the flagella, such that the cell can change its swimming direction, but in those cases light is sensed by a green light photosensor (Nagel 2002).

The most striking difference between our results in *S. pyriformis* and these prior studies of *P. bursaria* is that, in *P. bursaria*, the photosynthesis-dependent accumulation of cells in regions of bright light does not occur by directed swimming, but by a “kinetic mechanism” (Schnitzer 1990), in which the swimming speed is reduced whenever cells enter a region of higher light intensity, such that cells entering a region of brighter light spend more time there, eventually accumulating in the brightest areas. This effect is mediated by the generation of oxygen by photosynthesis which then regulates ciliary beating (Iwatsuki and Naitoh 1988; Cronkite 1981). Similar mechanisms have been reported in other ciliates (Finlay 1987).

In contrast, we have found that *S. pyriformis* swims directly toward the light source, a behavior quite different from the kinetic mechanism of photo-accumulation. One interesting possibility is that local oxygen production by the surface-docked algae could be influencing local ciliary motion in a manner similar to what happens on a whole-cell level in *P. bursaria*. Because of the larger size of *S. pyriformis*, it could be the case that increased oxygen caused by brighter light would cause slower motion of only the cilia near the illuminated part of the cell, causing the cell to steer towards brighter light. This model would require the cilia to respond to oxygen level variations generated by the algae on a time scale comparable to the rotational period of the *Stentor* cell (which is on the order of 1 second). Such a mechanism would require algae to be located near the cilia, which they are (**Figure 1**), but would not require the algal cells to be rotationally oriented, which is consistent with our electron micrographs that do not show any preferential orientation of the algal cells with respect either to the cell surface or the long-axis of the cell.

Given that the photosensor is inside the algal endosymbionts, but the motile apparatus is clearly the cilia of the host *Stentor* cell, we conclude that phototaxis in this organism thus appears to involve an exchange of information from symbiont to host. It is well known that symbiosis involves back and forth exchange of metabolic products. Phototaxis in *S. pyriformis* supports the idea that the exchange also involves information.

We note that our results only show that the alga and photosynthesis are needed, but this could be a permissive rather than instructive cue. We therefore cannot rule out a model in which the endosymbiont is a permissive cue required to allow phototaxis, but the actual photoreceptor to provide directional information is located in the host cell.

### Significance of surface association and microtubule baskets

The location of *Chlorella* cells at the surface of the *S. pyriformis* cell is quite striking, particularly so given the large size of the cells which leaves plenty of room in the interior to accommodate algal cells, as demonstrated by our centrifugation experiments. The retention of the endosymbionts near the surface appears to entail the development of a dense microtubule basket system, not seen in other *Stentor* species, presumably for the primary purpose of symbiont positioning. Could such location on the cell surface confer a fitness benefit to the host or to the symbionts?

The fact that *S. pyriformis* cells rotate while they swim indicates that each algal cell will spend part of the time fully exposed to incoming light and not blocked by any other symbionts, which would not be the case if the symbionts were filling up the interior of the cell. This could potentially help the *Chlorella* cells harvest more light than if they were distributed through the 3D interior of the cell.

Another possible physiological function of surface-docked symbionts could be to provide photoprotection for the host nucleus. By arranging the algae in a shell near the cell surface, with the macronucleus arranged as a small number of round nodes located in the cell interior, photoprotection of the host DNA by NPQ in the algae could be more efficient than if the algae and macronucleus were interspersed with each other, or if the macronucleus was located near the cell surface as it is in other *Stentor* species such as *S. coeruleus*. We note that the arrangement of the alga resembles the arrangement of melanin granules into “umbrellas” that partially surround the nucleus in human skin cells in such a way as to block incoming light (Kobayashi 1998).

Another possible way in which surface docking might be adaptive is that, as discussed above, the association of *Chlorella* cells with the surface near the cilia might help them signal to the cilia to control the direction of cell swimming for phototaxis. Finally, docking near the surface may simply aid with gas exchange between the symbionts and the water outside the host, an effect that seems to explain the docking of mitochondria near the surface of large ciliate cells (Fenchel 2014).

What cellular adaptations are required for the host cell to be able to locate the endosymbionts near the cell cortex? The extensive meshwork of microtubules at the cell cortex in *S. pyriformis* is not seen in non-symbiotic *Stentor* species such as *S. coeruleus*, which suggests it may be involved in holding the algae in place. In *P. bursaria*, microtubules play a role in cytoplasmic streaming (Nishihara 1999), which in turn is required for maximal algal proliferation (Takahashi 2007). The organization of microtubules into a cortical network in *S. pyriformis* suggests a different role, namely in anchoring the algae rather than moving them around. It will be interesting to determine whether other *Stentor* species that contain endosymbionts also display a similar microtubule organization.

### Stentor pyriformis as a model system for unicellular endosymbiosis

Given that much is already known about the cell biology of *Chlorella* endosymbiosis within *P. bursaria*, it is appropriate to ask what we can learn from using *S. pyriformis* as a model system. First, as a ciliate that is highly diverged from *Paramecium* with many unusual features, including a standard genetic code and extremely large cell size, *S. pyriformis* provides a useful comparison. Comparison between cell biological features of endosymbiosis in these two highly diverged ciliates will be especially important if they acquired *Chlorella* endosymbiosis independently, as seems likely to be the case.

The large size of *S. pyriformis* also offers opportunities to learn more about how mixotrophy contributes to aquatic food-webs. One of the interesting aspects of mixotrophy is that the host cell can obtain nutrients both from primary production within its endosymbionts and also from predation of other free-living predatory protists, thus tying together two distinct trophic levels. Because of its enormous size, *S. pyriformis* may be able to ingest larger prey compared to smaller mixotrophic ciliates, including other ciliates, colonies of algae or choanoflagellates, or even small animals like tardigrades. Thus this organism takes the crossing of trophic levels to an even greater extreme.

From a technical perspective, the large size and robust wound-healing and regeneration abilities of *Stentor* cells permit micromanipulation experiments that would be difficult in smaller cell types. In particular, the ability to bisect cells after centrifuging all the algae to one side may provide opportunities to study the effect of acute removal of endosymbionts, compared to the more gradual removal that can be achieved in other organisms such as *P. bursaria* using DCMU to inhibit photosynthesis (Reisser 1976) or cycloheximide to inhibit protein synthesis (Weis 1984). Because the loss of endosymbionts is gradual in both of those treatments, it may give the host cell time to adapt in a way that it would not be able to do under our acute surgical removal approach, and thus the two types of clearing strategies may provide different types of information about the host-symbiont interaction. Another consequence of the huge size of *S. pyriformis*, which is comparable to the size of leaves in some plants, is that it is so large that PAM fluorometry can be applied to single cells.

Finally, the genus *Stentor* is a classical system for studying cellular morphogenesis, but to date almost all experimental work has been done in just one species, *S. coeruleus*. Further development of *S. pyriformis* will potentially enable an “evo-devo” approach to the study of pattern formation and regeneration in single cells.

## Materials and Methods

### Collection and culturing of S. pyriformis

*S. pyriformis* cells were collected from a pond in Falmouth, MA, by collecting water from near the pond surface, along with submerged plant matter, and then manually picking *S. pyriformis* cells from the water under a dissecting microscope using a pipette. *S. pyriformis* cells were cultured following the same procedure used for culturing *S. coeruleus* (Lin 2018), using filtered pond water (Carolina Biological) and feeding the *Stentor* cells with *Chlamydomonas* cells grown in TAP medium (Harris 1989). *S. pyriformis* cells appear to be more sensitive to contamination than *S. coeruleus*, and it proved necessary to remove waste products from the cultures every few days.

### Microscopy of Stentor cells

For immunofluorescence, antibodies were diluted in antibody dilution buffer (AbDil: 2% BSA in PBS with 0.1% Triton X-100) and stored at 4°C. Cells were picked using a pipette with a cut-off tip and added to fresh MSM (Lin 2018) which was then removed and replaced with fresh MSM. Cells were then starved 12-24 hours (overnight) to minimize food vacuoles. The next day, cells were fixed by adding 500 µl ice cold methanol and incubated for 30 min at –20°C. Excess methanol was then removed by pipetting, followed by incubation in 500 µl of 1:1 PBS:Methanol for 5 min at room temperature, and then in 500 µl PBS for 10 min at room temperature. Cells were then blocked in 500 µl RT AbDil for 1 hour at room temperature. Primary antibody (anti β-tubulin primary antibody t0198 from Sigma) was added at a dilution of 1:500 in 250 µl of AbDil and incubated for 1 hour at room temperature.

After removing excess liquid from the pellet of cells, cell were washed 3 x 5min in 500 µl of PBS. Next, 250 µl of secondary antibody diluted 1:500 in AbDil were added along with DAPI diluted 10,000x, and incubated in the dark at room temperature for 1 hour. Excess liquid was then removed and cells washed 3x 5 min in 500 µl of PBS in the dark, then mounted on a slide and coverslip using Vectashield media.

For electron microscopy, *S. pyriformis* cells were gently pelleted and fixed for transmission electron microscopy with 2.5% glutaraldehyde in 0.1 M sodium cacodylate buffer, pH 7.4, for one hour at room temperature. After three five-minute buffer washes, cells were post-fixed with 1% buffered osmium tetroxide for one hour, washed in deionized water and dehydrated through increasing concentrations of ethanol (30%, 50%, 75%, 90%, 100%, 100%, ten minutes each). Following two changes in propylene oxide (10 minutes each), the samples were infiltrated in a 1:1 mixture of propylene oxide/Polybed 812 epoxy resin (Polysciences, Inc., Warrington, PA) for three hours. The pellets were infiltrated overnight in 100% Polybed 812 epoxy resin followed by embedment in fresh epoxy resin and polymerization at 60°C for 24 hours. Resin blocks were trimmed to approximately 2 mm^2^ and sectioned at 70 nm using a Leica Ultracut UCT ultramicrotome (Leica Microsystems, Inc., Buffalo Grove, IL) and a Diatome diamond knife (Electron Microscopy Sciences, Fort Washington, PA). Ultrathin sections were mounted on 200 mesh copper grids and stained with 4% aqueous uranyl acetate for 10 minutes and Reynold’s lead citrate for 8 minutes to enhance contrast (Reynolds, 1963). Samples were observed using a JEOL JEM-1230 transmission electron microscope operating at 80 kV (JEOL USA INC., Peabody, MA) and digital images were taken using a Gatan Orius SC1000 CCD camera with Gatan Microscopy Suite version 3.10.1002.0 software (Gatan, Inc., Pleasanton, CA).

### Expansion Microscopy

We used the U-ExM expansion microscopy protocol (Gambarotto 2019). *S. pyriformis* cells were adhered to poly-lysine coated coverslips for 7 min at room temperature, pre-fixed in 3% formaldehyde in PBS for 10 min at RT, washed once in PBS, then fixed in 0.7% formaldehyde with 0.15% or 1% acrylamide in PBS for 4–5 h (with agitation).

5 µL TEMED and then 5 µL APS were added to 90 µL of MS (19% sodium acrylate (SA), 10% acrylamide (AA – 40%), 0.1% N,Nʹ-methylenebisacrylamide (BIS – 2%) in 1x PBS to yield final concentrations of 0.5% TEMED and 0.5% APS. 35 µL of this mix was then added to each 12 mm coverslip on parafilm

Polymerization was started at 4°C for 5 min, and then the samples were incubated at 37°C in the dark for 1 h. The gel was then placed in ∼2 ml of denaturation buffer (200 mM SDS, 100 mM NaCl, 50 mM Tris in ultrapure water, pH 9 2.88 g SDS, 0.584 g NaCl, 0.303 g Tris in 50 mL MilliQ H2O, pH 9 with 1 M HCl) in a six-well plate and Incubated for 15 min at RT with agitation. Gels were then removed from the coverslips with flat tweezers, moved into a 2 ml Eppendorf centrifuge tube filled with fresh denaturation buffer, and incubated at 95°C for 30 min. After denaturation, gels were placed in beakers filled with ddH2O for the first expansion. Water was exchanged at least twice every 30 min at RT, and then gels were incubated overnight in ddH2O.

Excess water was then removed by placing the gels in PBS two times for 15 min. Note that in this step, gels shrank back to ∼50% of their expanded size. Gels were then incubated with primary antibody diluted in 2% PBS/BSA at RT for ∼3 h, with gentle shaking.

Gels were then washed in PBS-T three times for 10 min with shaking and subsequently incubated with secondary antibody solution diluted in 2% PBS/BSA for ∼3 h at RT with gentle shaking. Gels were then washed in PBS-T three times for 10 min with shaking and finally placed in beakers filled with ddH2O for expansion. Water was exchanged at least twice every 30 min, and then gels were incubated in ddH2O overnight. We found that the gels expanded between 4.0× and 4.5× according to SA purity.

Gels were then mounted as follows. Each gel was placed in a 10 cm dish and excess water removed with a kimwipe. A small piece of the gel was then cut and transferred smooth side down on a PDL coated 35 mm glass bottom dish. The gel was then immobilized with 1% LMA and imaged with a 60x n.a. 1.3 oil lens, using a z-step size of 1 micron.

### Isolating Chlorella from Stentor pyriformis

Approximately 300 *S. pyriformis* cells were collected into a minimal volume of filtered pond water (2 mL total), sonicated until most cells were lysed, and then 500 μL of lysate were layered on top of 1 mL of 60% Percoll and centrifuged at 8k rpm for 1 min. After removing supernatant, the green algal pellet was resuspended in 1 mL of MBBM, centrifuged at 4k rpm for 1 min, and then resuspended in MBBM. *Chlorella* cells were subsequently cultured in MBBM media, both in liquid cultures and on 0.8% agar plates.

### Phylogenetic analysis of S. pyriformis host and endosymbionts

DNA isolated from *S. pyriformis* cells was used to perform PCR using the Thermo Phire plant direct PCR kit with the following primers:

For *S. pyriformis*: Universal Primers ∼ 500bp amplicon (Tm: 59°C)

VB017_18S rRNA_F (CG167_18S rRNA_F): CAG CAG CCG CGG TAA TTC C

VB018_18S rRNA_R (CG168_18S rRNA_R): CCC GTG TTG AGT CAA ATT AAG C

For algae in *Stentor*: ∼ 350bp amplicon (Tm: 65°C)

VB001_18S_rRNA_Pyriformis_F: AAATTAGAGTGTTCAAAGCAGGC

VB002_18S_rRNA_Pyriformis_R: GTCTGGACCTGGTAAGTTTCCC

For algal *rbcL* sequence (Tm: 61°C)

VB003_RbcL_F: TTACGTTTAGAAGATCTTCGTATTCCAC

VB004_RbcL_R: GCTAATTCAGGACTCCATTTGCA

16S rRNA sequencing (Tm: 55°C)

VB015_16s_27f_F: GAGAGTTTGATCCTGGCTCAG

VB016_16s_1492r_R: GGTTACCTTGTTACGACTT

DNA sequencing was performed using 200 ng of each PCR product. We generated fasta files containing our 18S or rbcL sequences as well as sequences from related species found on NCBI databases. We performed multiple sequence alignment using a ClustalW (Thompson 1994) web server (https://www.genome.jp/tools-bin/clustalw) and trimmed alignments with gblocks (Talavera 2007). We then used the same webserver to run PhyML (Guindon 2010) with 100 bootstraps and used FigTree (https://github.com/rambaut/figtree/releases/tag/v1.4.4) for visualization.

### PAM measurement of photosynthetic efficiency of Chlorella inside the host cell

PAM chlorophyll fluorescence measurements were performed using the Dual-PAM-100 (Heinz Walz) on either *S. pyriformis* or cultured *Chlorella* cells as previously described (Brooks and Niyogi., 2011). *S. pyriformis* and *Chlorella* cell PAM measurements were performed using 500 µmol photons m^-2^ s^-1^ actinic light and 986 µmol photons m^-2^ s^-1^ of saturating light. *Chlorella* PAM measurements were all performed at 5.5 x 10^5^ cells/mL under conditions indicated in the figure legends. *S. pyriformis* PAM measurements were performed using ∼300 cells isolated from Morse pond within 24 hours, suspended in 2 mL of filtered pond water.

### Apparatus for phototaxis measurement

The apparatus in Figure 6A was constructed using LEDs, LED drivers, and LED power supplies from Thorlabs, with bandpass filters to select illumination wavelength and an aspheric condenser lens held with a lens tube (Thorlabs). Images were acquired using a Raspberry Pi HQ camera (16 mm, 5 Mpix) with a CS lens, controlled by a Raspberry Pi 4 desktop kit (4Gb).

### Chlorella Genomic DNA extraction

DNA was extracted from *Chlorella* using a procedure modified from methods used for the alga Nannochloropsis (Radakovits 2012; Vieler 2012; Gee 2017). *Chlorella* cells were grown in 50 mL MBBM to mid-log phase (approximately 5×10 ^6^ to 2×10^7^ cells/mL). in a 125 mL sterilized Erlenmeyer flask shaking at ∼100 rpm in continuous 100 µmol photons m^-2^ s^-1^ light at constant 28 °C. Cells were collected by transferring to 50 mL conical tubes and centrifuging in benchtop centrifuge (1000 rcf) for 5 min, resuspended in 10 mL ddH2O to rinse away media, and re-pelleted by centrifugation. Cells were lysed by adding 800 µL of 1x CTAB lysis buffer (2% cetyltrimethylammonium bromide, 100 mM Tris-HCl pH 8.0, 1.4 M NaCl, 20 mM EDTA) with 0.8 µL of 100 µg/mL proteinase K (stock solution containing 50 mM Tris (pH 8.0) and 10 mM CaCl2) to the pellet in 2mL round-bottom microcentrifuge tubes (MCT), and incubate at 50 °C for 30 min to lyse cells and digest proteins. The lysate was then phenol/chloroform extracted using 3.5 mL of 1:1 buffered (pH ∼7.5-8.0) phenol and chloroform mixture and centrifuged at 20,000 rcf) for 2.5 min. The upper, clear, colorless aqueous phase was transfer to new MCT. RNase A was added to a concentration of 100 µg/mL (8 µL of 10 mg/mL stock) and incubated at 37°C for 60 min. If RNA contamination is found to be a problem, increase concentration and incubation. The phenol/chloroform extraction was then repeated, transferring to another clean MCT.

Two final extractions were then performed with 100% chloroform to remove trace phenol using same volumes and spins. After last extraction, 550 µL was transferred to fresh MCT to avoid collecting the interface. DNA was then ethanol precipitated by adding 2.5 volumes (e.g. 1.375 mL) of ice-cold 100% EtOH and 0.1 volume of 3M sodium acetate (e.g. 55 µL). After precipitation for 60 min at –20 °C, tubes were centrifuged at 4°C for 15 min at 20,000 g. The pellet was washed with 1 mL ice-cold 70% EtOH and centrifuged again for 5 min max speed at 4°C to make sure pellet is seated securely again.

The pellet was then dried in a stream of filtered air, and resuspended in 30 µL of water. DNA quality was checked by Nanodrop spectrophotometry followed by a fluorometric Qubit assay.

### Sequencing of the Stentor pyriformis and Chlorella variabilis genomes

Total genomic DNA samples from *Stentor pyriformis* were sequenced using an Oxford Nanopore MinION, R9.4 flow cell, and ligation sequencing prep (LSK109), which produced 0.5 million reads with an N50 size of 6 kbp and a total of 1.7 Gbp of sequence. Nanopore sequencing reads were assembled using Flye (Kolmogorov 2019) followed by polishing with Medaka (nanoporetech.com).

No algal reads were obtained during the sequencing and assembly process using DNA obtained from *S. pyriformis* cells, which is to be expected given the difficulty of breaking algal cell walls. For this reason, DNA was isolated from *Chlorella* cells obtained from *S. pyriformis* and cultured on MBBM plates. Sequencing and assembly of the *Chlorella* genome was performed the same as for *S. pyriformis*.

Nanopore sequencing genomic DNA from Stentor pyriformis and its endosymbiont, Chlorella variabilis will be available at Data Dryad. https://doi.org/10.5061/dryad.g79cnp612 The genome assembly for both *S. pyriformis* and its *Chlorella* endosymbiont presented in this paper are available online at StentorDB (Stentor.cliate.org).

### RNA-seq and Gene prediction for Stentor pyriformis

Full code and descriptions for RNA-Seq alignment and gene prediction, including data on intron lengths and contig sizes, are available at https://github.com/aralbright/2022_pyriformis. We altered the source code of HISAT2 (Kim 2019) to accommodate for short introns as found in other heterotrichs (Singh et al 2023), and the closely related *S. coeruleus* (Slabodnick et al 2017). The variable minIntronLen in hisat2.cpp was reduced to 9. HISAT2 with this change was run with the following parameters: –-min-intron-len 9 –-max-intron-len 101 –-very-sensitive. We used samtools (Li et al 2009) to sort and index the alignments.

Genes were predicted using Intronarrator (https://github.com/Swart-lab/Intronarrator), which predicts and removes introns before running Augustus with an intronless model and adds back the introns at the end. We ran Intronarrator by altering the path variables in intronarrator.sh as well as the following parameters: MAX_INTRON_LEN=101 and GENETIC_CODE=1. To standardize the gene names, we modified a GFF Parser (Hannon and Eisen 2024) and added custom scripts to change the names of contigs to StePyr_contig_# and genes to StePyr_#.

## Acknowledgments

We thank Ben Jenkins for many helpful suggestions and comments about ciliate endosymbiosis, and Brandon Seah for insightful comments on the manuscript. This work was funded by the Gordon and Betty Moore Foundation Symbiosis in Aquatic Systems Initiative SMS Grant GBMF9348 (WFM), as well as by the Howard Hughes Medical Institute (KKN), NIH grant R35 GM130327 (WFM), 5K12GM081266 (ARA), and an MBL Whitman Fellowship (VB). The Microscopy Services Laboratory, Department of Pathology and Laboratory Medicine, is supported in part by P30 CA016086 Cancer Center Core Support Grant to the UNC Lineberger Comprehensive Cancer Center. KKN is an investigator of the Howard Hughes Medical Institute. This article is subject to HHMI’s Open Access to Publications policy. HHMI lab heads have previously granted a nonexclusive CC BY 4.0 license to the public and a sublicensable license to HHMI in their research articles. Pursuant to those licenses, the author-accepted manuscript of this article can be made freely available under a CC BY 4.0 license immediately upon publication.

## List of Supplemental Materials

**Supplemental Video S1**. Live Stentor pyriformis cells imaged under transmitted light.

**Supplemental Video S2**. Expansion microscopy of S. pyriformis cells stained with anti-tubulin antibodies, stepping in the z axis in 1 micron increments starting at the surface of the cell and moving inwards.

**Supplemental Video S3**. Time lapse fluorescence imaging shows *Chlorella* endosymbionts changing position on the surface of *S. pyriformis*.

**Supplemental Video S4**. *S. pyriformis* cells imaged in the phototaxis apparatus of Figure 6A. Light source is located to the left.

**Supplemental Figure S1**. The *Chlorella variabilis* genome contains genes encoding key photoprotection proteins involved in NPQ.

## Notes

### Competing Interest Statement

The authors have declared no competing interest.

