## Supplementary figures and images for "The cell biology and genome of *Stentor pyriformis*, a giant cell that embeds symbiotic algae in a microtubule meshwork"

### Supplemental Figure S1

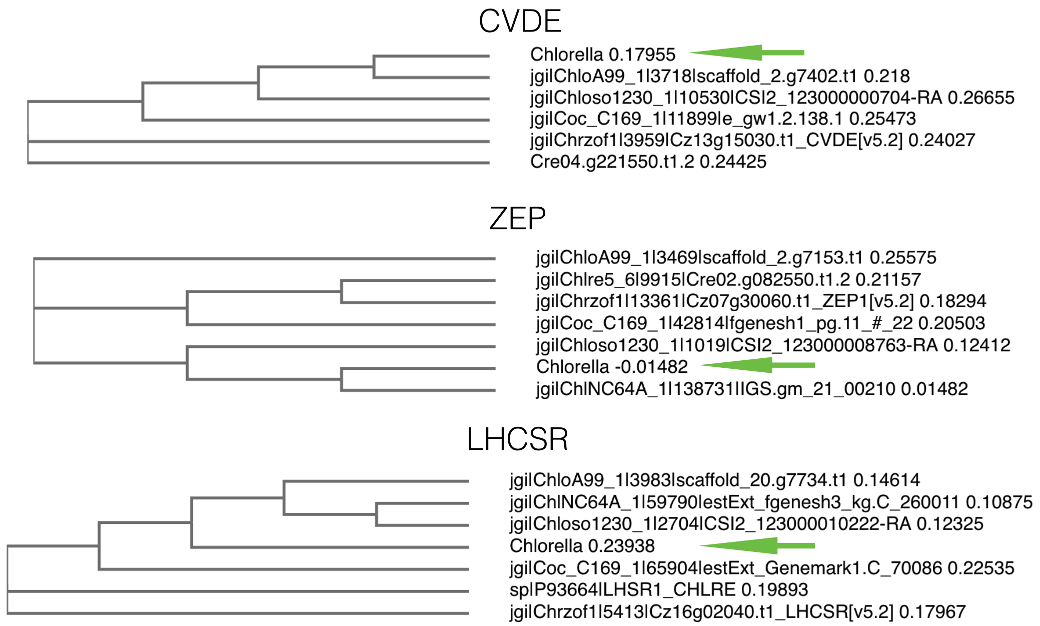


**Supplemenal Figure S1**
